# State Dependent Anionic Pore Currents Conducted by Single Countercharge Mutants in a Voltage-Sensing Phosphatase

**DOI:** 10.1101/2022.04.04.487073

**Authors:** Rong Shen, Benoît Roux, Eduardo Perozo

## Abstract

Mutating gating charge residues in the S4 segment of voltage-sensing domains (VSDs) can cause ionic leak currents through the VSDs. These leak currents, known as gating pore or omega currents, play important pathophysiological role in many diseases. Here, we show that mutations in a key countercharge residue, D129, in the *Ciona intestinalis* voltage-sensing phosphatase (Ci-VSP) facilitate conduction of unique anionic omega currents. Neutralization of D129 causes a dramatic positive shift of activation, facilitates the formation of a continuous water path through the intermediate state VSD, and creates a positive electrostatic potential landscape inside the VSD leading to anion selectivity. Increasing the population or duration of the conducting state by a high external pH or an engineered Cd^2+^ bridge markedly increases the current magnitude. Our findings uncover a new role of countercharge residues and could inform on the mechanisms of channelopathies linked to countercharge residue mutations.

## Introduction

Canonical voltage-sensing domains (VSDs) comprise four transmembrane segments S1-S4, forming an antiparallel helix bundle. VSDs undergo defined conformational transitions in response to membrane potential changes to orchestrate the gating of the pore domain of ion channels(*1*–*4*), the catalytic activity of phosphatases (*5*–*8*), the activation of ion exchange of solute carriers (*9*, *10*), and the proton-specific conduction of proton channels (*11*–*17*). VSDs share high sequence and structural similarity with each other, particularly the conserved positive gating charge residues on S4 and negative countercharge residues on S1-S3 (*18*–*20*), and a layer of in-plane hydrophobic residues on S1-S3 (*21*–*24*), thought to electrically separate the extracellular and intracellular aqueous crevices of the VSDs, preventing passage of water and ions and ensuring a focused electric field (*24*–*27*). The number and location of these gating charges and countercharges vary in different VSDs, resulting in diverse voltage gating set point and sensitivity (*28*–*31*).

The omega current has been identified as an alternative pathway for ion translocation, distinct from the pore currents evoked by the opening of the central pore domain or the transient gating currents (*25*). Instead, omega currents permeate through the VSDs themselves, displaying proton and/or cation selectivity. In *Shaker* potassium channels, mutating the first S4 gating arginine to histidine can create steady proton currents at hyperpolarization potentials (*25*). Subsequent studies showed that neutralization of a single, two or three adjacent gating charge residues in S4 of potassium channels (*32*–*37*), as well as sodium and calcium channels (*38*–*44*), can generate a monovalent cation selective current through the VSDs at depolarizing or hyperpolarizing potentials reaching as much as 1% of the central pore current magnitude (*32*, *43*). Large organic monovalent cations tetraethylammonium (TEA^+^) and N-methyl-D-glucamine (NMDG^+^) are much less permeable and the monovalent cation currents can be blocked by divalent cations at millimolar concentrations (*39*). Cationic omega currents have also been observed at hyperpolarized potentials in a truncated wild type (WT) VSD of *Shaker* potassium channel (*45*), and the WT VSD in the α3 subunit of the ascidian CatSper calcium channel (Ci-CatSper3) (*46*), in both truncated and full-length forms. The WT VSD of Ci-CatSper3 can even conduct divalent cations Ca^2+^, Ba^2+^ and Sr^2+^ (*46*). Although voltage-sensitive phosphatases can neither conduct ions nor permeate protons like proton channels, mutations of S4 gating charge residues in Ci-VSP have been shown to conduct protons as well (*47*, *48*). The importance of role of these cationic omega currents caused by missense mutations is highlighted in certain inherited channelopathies, e.g., hypokalemic periodic paralysis, normokalemic periodic paralysis and cardiac arrhythmias, and may also have been linked to other pathologies (*35*, *39*, *40*, *43*, *44*, *49*–*52*).

As the direct sensing component for membrane potential, gating charge residues in S4 of VSDs have attracted a great of attention, however, the physiological and pathological functions of missense mutations of countercharge residues continue to be understudied (*32*, *53*). A fluorescent voltage sensor derived from the VSD of Ci-VSP (Ci-VSD) containing a countercharge residue mutation displays omega currents, but the nature of the charge carrier remains undetermined (*54*). We have explored the role of countercharge residues in voltage sensitivity and their role in generating omega currents by focusing on residues D129, D136 and D151 at the extracellular side of Ci-VSD. We demonstrate here that single neutralizing mutations at D129 in S1 lead to anionic omega currents through the intermediate “Down” state VSD. These mutations cause large positive shifts in the voltage dependent of activation, further populating the “Down” state and amplifying the influence of external pH on activation gating. These effects drive a steady anion current upon activation and a transient anion current upon deactivation, which can be enhanced by a engineered Cd^2+^ bridges between S4 and lipids (immobilizating S4 during deactivation). In agreement with these observations, MD simulations and free energy calculations show that neutralization of D129 or R226 generate significant changes in the electrostatic potential landscape inside the gating pore. In turn, these catalyze the anionic currents observed in the D129 mutants and the cationic currents in the R226 mutants.

## Results

### External pH modulates anionic omega current in the D129A mutant of Ci-VSP

Voltage-sensing phosphatases contain a N-terminal transmembrane VSD and a C-terminal cytosolic membrane-targeted enzyme domain. The ascidian VSP, Ci-VSP, was the first VSP that exhibited both voltage sensitivity and catalytic activity (*5*). Phylogenetic analysis shows that the VSDs of Voltage-sensing phosphatases are closer to the VSDs of voltage-gated proton channels (Hv1s) than those of voltage-gated ion channels (*15*, *55*). A sequence alignment shows that both VSPs and Hv1s have a conserved countercharge aspartate in S1 (Fig. 1, A and B), a helix-turn above the hydrophobic gasket residues (*21*). This residue (D112) has been reported to play a role as hHv1 selectivity filter, as neutralizing D112 generates anion currents (*55*). In Ci-VSP an asparagine substitution at this position (D129N) results in a omega current at positive voltages (*54*).

**Fig. 1.**
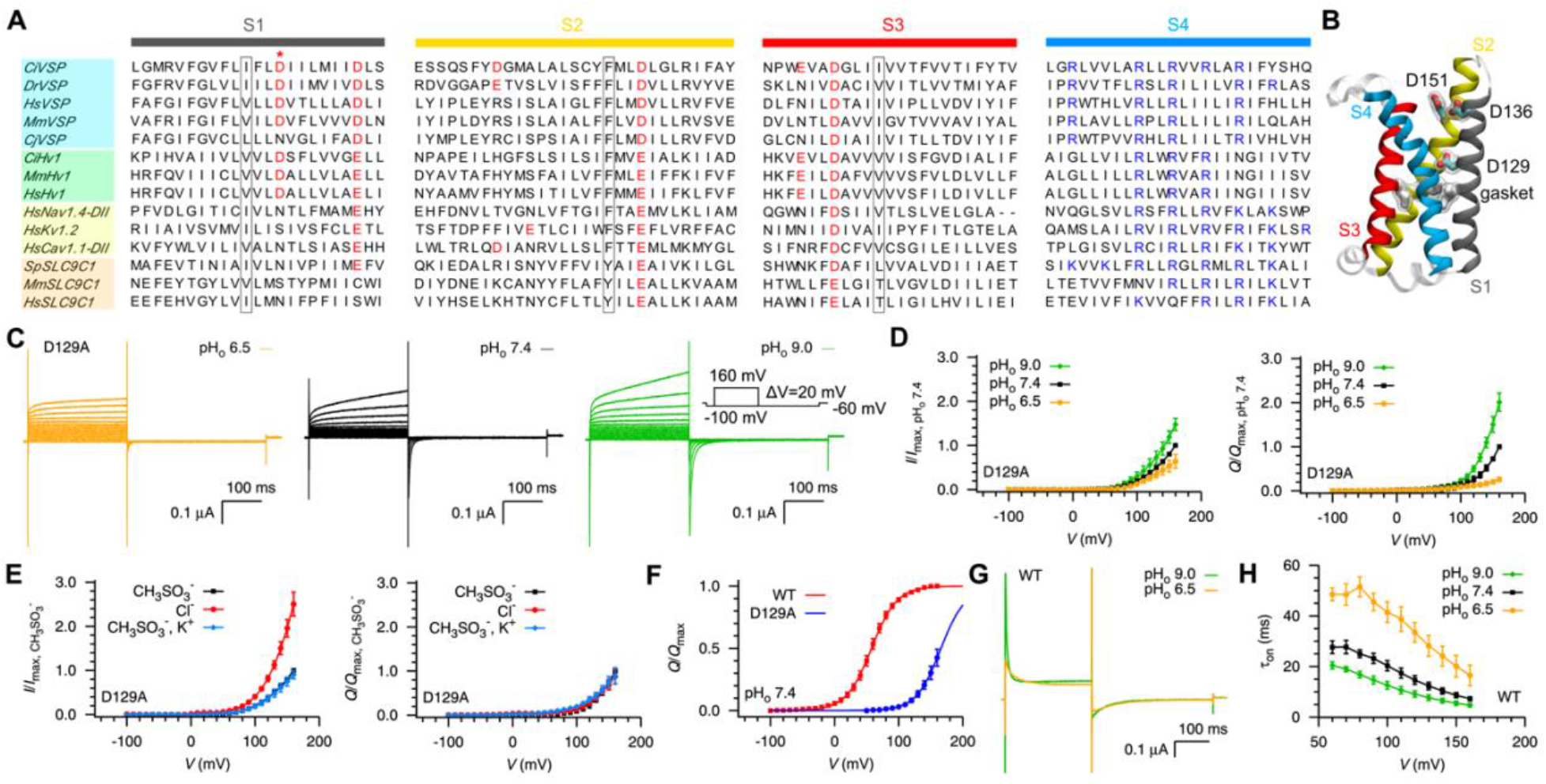
Anionic currents elicited by mutating countercharge residue D129 in Ci-VSP. (**A**) Sequence alignment of the transmembrane segments S1-S4 of representative VSDs of voltage-sensing phosphatases, proton channels, voltage-gated ion channels and sperm-specific solute carriers SLC9C1 from *Ciona intestinalis* (*Ci*), *Danio rerio* (*Dr*), *Homo sapiens* (*Hs*), *Mus musculus* (*Mm*), *Callithrix jacchus* (Cj), *Strongylocentrotus purpuratus* (*Sp*). Negative countercharge residues in S1-S3 and positive gating charge residues in S4 are colored in red and blue, respectively. Hydrophobic gasket residues in S1-S3 are highlighted in black rectangles. Red star indicates the D129 site. (**B**) Crystal structure of Ci-VSD in the “Up” state (PDB: 4g7v). The hydrophobic gasket residues: I126 (in S1), F161 (in S2), I190 (in S3); and countercharge residues facing the extracellular crevice: D129 and D136 (in S1), D151 (in S2) are highlighted in stick and transparent surface representation. (**C**) Representative currents for D129A mutant of Ci-VSP at different pH values (pH_o_) of the external recording buffer containing 120 mM NMDG^+^/CH_3_SO_3_^-^. Inset is the voltage pulse protocol used to elicit the currents. (**D**) Normalized *I-V* (left) and *Q-V* (right) curves for D129A mutant of Ci-VSP with respect to the corresponding maximum value at pH_o_ 7.4, where *I* is the amplitude of ON current at the end the pulse after linear leak subtraction and *Q* was the net translated charge calculated by integrating OFF current. Error bars are standard deviation (*n* = 4, 12, 7 for pH_o_ 6.5, pH_o_ 7.4, pH_o_ 9.0, respectively). The external recording buffer contains NMDG^+^/CH_3_SO_3_^-^ as in (C). (**E**) Normalized *I-V* (left) and *Q*-*V* (right) curves for D129A mutant of Ci-VSP at different external recording buffer conditions, containing 120 mM NMDG7CH_3_SO_3_^-^ (CH_3_SO_3_^-^), 120 mM NMDG^+^/Cl^-^ (Cl^-^) or 108 mM NMDG^+^ + 12 mM K^+^/CH_3_SO_3_^-^(CH_3_SO_3_^-^, K^+^). The data were normalized with respect to the corresponding maximum value at the CH_3_SO_3_^-^ condition. Error bars are standard deviation (*n* = 4, 3, 12 for Cl^-^, CH_3_SO_3_^-^, K^+^, CH_3_SO_3_^-^, respectively). (**F**) Non-linear least-squares fitting of the *Q*-*V* curves for wild type (WT) and D129A mutant of Ci-VSP at pH_o_ 7.4 with the external recording buffer of 120 mM NMDG^+^/CH_3_SO_3_^-^. (**G**) Representative current of WT Ci-VSP at the depolarizing voltage pulse of 160 mV, recorded at pHo 6.5 or 9.0 with the external recording buffer of 120 mM NMDG^+^/CH_3_SO_3_^-^. (H) Activation time constant for WT Ci-VSP at different pHo with the external recording buffer of 120 mM NMDG^+^/CH_3_SO_3_^-^.

In order to understand the functional role of D129 in Ci-VSP voltage gating, we electrophysiologically evaluated the alanine substitution of D129 experiments using the cut-open oocyte voltage-clamp technique (*56*, *57*). D129A is characterized by abnormal gating currents in non-permeant 120 mM N-methyl-D-glucamine (NMDG^+^) methanesulfonate (CH3SO3^-^) at various pH conditions (Fig. 1, C and D): (1) it activates slowly without a typical gating peak at the beginning of the ON gating currents (*5*); (2) it conducts an outward current at depolarization potentials; (3) the magnitude of the ON current at the end of the depolarization pulse (*I*) and the net OFF gating charge (*Q*) are tightly regulated by external pH in the same manner. Since the external buffer contains only large organic ions (NMDG^+^ and CH_3_SO_3_^-^), we first suspected that the currents were carried out by protons or hydroxides. If the mutant conducts protons, inward currents should increase at low pH, as is the case for omega currents in Ci-VSP gating charge mutants of (*48*, *58*). However, inward currents decreased at low pH. If, on the other hand, the mutant is permeable to hydroxides, inward currents should decrease at high pH (outflow of hydroxide). Yet, currents increased at high pH (Fig. 1, C and D).

To define D129A charge carrier, we replaced the external cation from 120 mM NMDG^+^ with a combination of 108 mM NMDG^+^ and 12 mM K^+^ in one experiment and the external anion from CH3SO3^-^ completely to Cl^-^ in the other experiment (Fig. 1E) at a constant pH 7.4. The inclusion of K^+^ did not change the maximum *I* or *Q*. However, ON currents at 160 mV increased by two to three times after replacing CH_3_SO_3_^-^ with Cl^-^ without a substantial decrease in the *Q* or OFF currents. It should be noted that *Xenopus* oocytes have endogenous cation and chloride channels that generate outward currents at large positive potentials (*59*, *60*), but these endogenous channels do not generate OFF currents, are less influenced by external pH, and the maximum magnitude and increment of ON currents in the external Cl^-^ solution is smaller than what we see with the D129A mutant (fig. S1). Rather, the expectation was that the D129A mutant would conduct anion-selective omega currents regulated by pH, and that the omega current would be eliminated once the VSD deactivated. Furthermore, the rapid closure of the gating pore might have prevented conduction of any outward anion currents upon deactivation (the OFF currents include OFF gating currents with no tail currents) (*46*), or the small size of the omega currents in the D129A mutant (changes of the tail currents are negligible compared to the OFF gating currents) (*32*, *43*).

A large positive shift in gating voltage sensitivity has also been observed in the D129A mutant of Ci-VSP; its activation V_1/2_ shifting more than 100 mV relative to WT (Fig. 1F). We noticed that the activation kinetics of the WT and mutants of Ci-VSP can be tuned by external pH (Fig. 1, G and H), where higher pHs lead to faster gating kinetics. In other words, the transition rate between adjacent conformational states increased at high pH. However, there was no change in WT Ci-VSP net translocated charge (*Q*), and the *Q-V* curve reached a plateau at pHo between 6.5, and 9.0 (*48*). We argue that under these conditions the D129A mutant does not reach the “up-plus” state of the VSD (fully activated), even at +160 mV (*48*). At high pH, the transition of the D129A mutant from the non-conducting resting state (“down-minus” state), to the intermediate conducting state(s) accelerates, increasing the population of the conducting state(s).this is a consequence of the blockage of the further transition to the fully activated state which may or may not conduct anions.

### Augmenting of the anionic omega current by an engineered Cd^2+^ bridge

Given that the anionic omega current is state dependent, we pursued alternative ways to increase to increase the omega current (the lifetime of the conducting state(s)), while decoupling the pH modulation effect. We took advantage of the finding that a cysteine mutation at the extracellular side of S4 can form Cd^2+^ bridges with lipid headgroups, immobilizing S4 in its activated conformation (*48*). To that end, we introduced a cysteine at position V219 in the background of the D129A mutant (Fig. 2A). Control experiments showed that V219C does not show omega currents and external Cd^2+^ have little or no effect on D129A gating (fig. S2). The V219C/D129A double mutant displayed a similar behavior as the single D129A mutant (fig. S2D), and is able to conduct inward Cl^-^ currents upon activation (Fig. 2B).

**Fig. 2.**
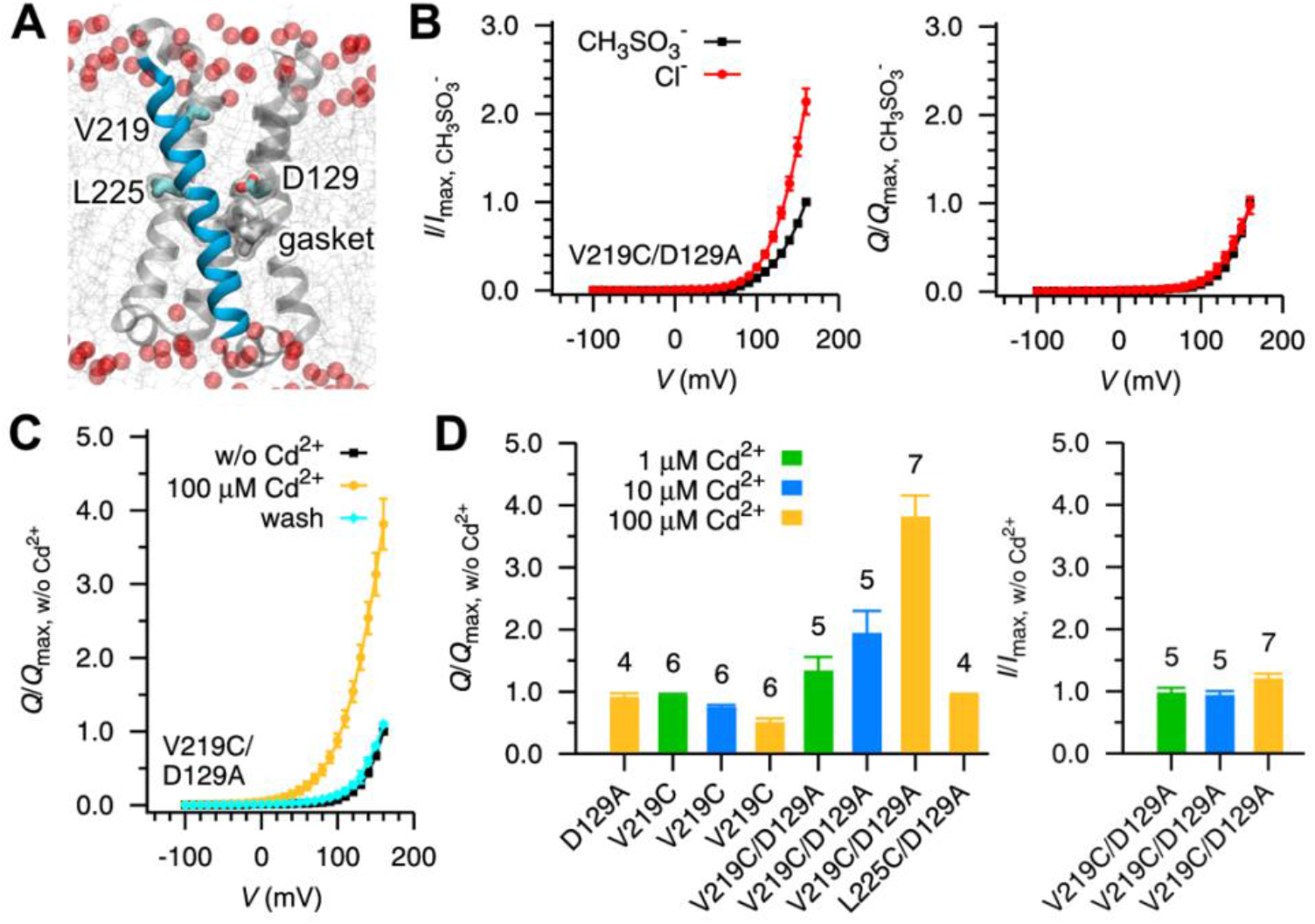
State dependent anion conduction enhanced by immobilizing the segment S4 via engineered Cd^2+^ bridges. (**A**) A snapshot from a MD simulation of the “Up” state Ci-VSD. D129 in S1, hydrophobic gasket residues, and V219 and L225 in S4 are shown in stick and transparent surface representation. For clarity, only parts of the lipid molecules and their non-ester phosphate oxygen atoms are shown in gray lines and transparent spheres, respectively. (**B**) Normalized *I-V* (left) and *Q*-*V* (right) curves for the double mutant V219C/D129A of Ci-VSP with external recording buffer of 120 mM NMDG^+^/CH_3_SO_3_^-^ (CH_3_SO_3_^-^) or 120 mM NMDG^+^/Cl^-^ (Cl^-^). The data were normalized with respect to the corresponding maximum value at the CH_3_SO_3_^-^ condition. Error bars are standard deviation (*n* = 5 for both conditions). (**C**) Normalized *Q*-*V* curves for the double mutant V219C/D129A of Ci-VSP in the absence (black), in the presence (yellow) and after washout (cyan) of 100 μM Cd^2+^. The data were normalized with respect to the corresponding maximum *Q* in the absence (w/o) of Cd^2+^. Error bars denote standard deviation (*n* = 7). (**D**) Cd^2+^ concentration dependent immobilization of S4 by the V219C mutant, but not L225C.The *Q*-*V* (left) and *I*-*V* (right) curves were normalized with respect to the corresponding maximum value in the absence (w/o) of Cd^2+^. Error bars denote standard deviation with the number of *n* being listed on the top of each bar.

Addition of external 100 μM Cd^2+^ leads to an increase in OFF currents for the V219C/D129A double mutant (fig. S2D and Fig. 2C), in contrast to the behavior of the V219C mutant (fig. S2B and Fig. 2D). The increase in OFF currents for the V219C/D129A double mutant can be attributed to outward anion currents, as the engineered Cd^2+^ bridge slows down the closure of the gating pore. As expected, higher the concentrations of Cd^2+^, lead to reductions in V219C net charge. Whereas for the V219C/D129A double mutant, Cd^2+^ -driven increases in net charge *Q* readily reverse by washout with the chelating buffer (Fig. 2, C and D). In comparison, the ON currents only show a slight increased even at 100 μM Cd^2+^ (Fig. 2D).

### Anionic omega currents vary in single D129 mutants

The identification of small but distinguishable anionic omega currents conducted by the D129A mutant encouraged us to make other uncharged amino acid substitutions at D129 (*32*, *55*). Aasparagine, leucine and valine substitution at D129 can conduct outward currents under positive potentials in NMDG^+^/CH_3_SO_3_^-^ buffer at neutral pH, displaying inward gating currents and/or tail currents upon deactivation (Fig. 3A and fig. S3). The magnitude of *I* and *Q* for each D129 mutant changes in the same direction (Fig. 3B), in contrast to the WT which yields a large net OFF charge *Q* but a small ON current *I* because of its impermeability of omega currents. Replacing the external anion CH_3_SO_3_^-^ with Cl^-^ increases the ON currents while decreases the OFF currents, consequence of the inhibition of Cl^-^ outward flow (Fig. 3, C and D and fig. S3, A to C). Notably, asparagine substitution of D129, a naturally occurring mutation in certain VSPs (Fig. 1A), has a about 5-fold increase of the maximum ON current at +160 mV in the Cl^-^ solution than in the CH_3_SO_3_^-^ solution. Since the D129V mutant led to the largest omega currents, we performed extra experiments on it with other solutions (fig. S3, D to F): Replacing a fraction of external NMDG^+^ with K^+^ has little or no effect on both the ON and OFF currents. Decreasing the external ionic strength by dilution of the external solution NMDG^+^/CH_3_SO_3_^-^ with 50% isotonic sucrose decreases the ON currents (about 30% at +160 mV) but has little effect on the OFF currents. Changing the external anion CH_3_SO_3_^-^ with glutamate (Glu^-^) significantly decreases the ON currents (about 80% at +160 mV) but has little effect on the OFF currents (Fig. 3, C and D). These results confirm the anion selectivity of the D129 neutralization mutants (with a higher permeability for Cl^-^ than CH_3_SO_3_^-^), and the presence of anionic omega currents during both activation and deactivation. Dilution of external CH_3_SO_3_^-^ and the substitution of external CH_3_SO_3_^-^ with Glu^-^ have little effect on the OFF currents, confirming that they are a mixture of gating currents and omega currents or tail currents carried by cytosolic Cl^-^.

**Fig. 3.**
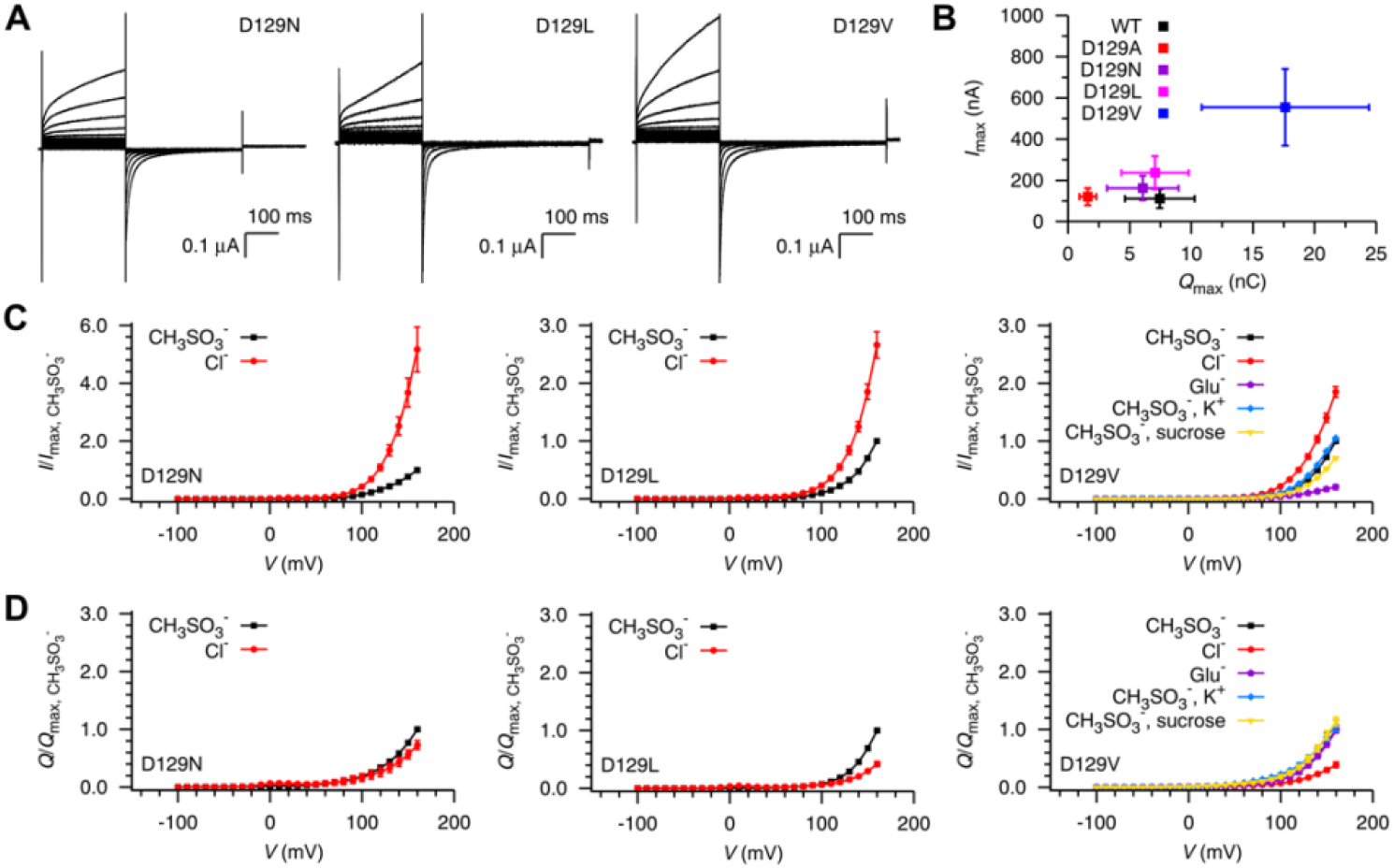
Anionic currents evoked in single D129 mutants. (**A**) Representative currents of D129N, D129L and D129V mutants of Ci-VSP with external recording buffer of 120 mM NMDG^+^/CH_3_SO_3_^-^. (**B**) *Imax* versus *Q*max of WT and D129 mutants of Ci-VSP recorded with external buffer of 120 mM NMDG^+^/CH_3_SO_3_^-^. Error bars are standard deviation (WT, *n* = 9; D129A, *n* = 8; D129N, *n* = 26; D129L, *n* = 9; D129V, *n* = 12). (**C**and **D**) Normalized *I-V* (C) and *Q*-*V* (D) curves for the D129 mutants of Ci-VSP with external recording buffer of 120 mM NMDG7CH_3_SO_3_^-^ (CH_3_SO_3_^-^), 120 mM NMDG^+^/Cl^-^ (Cl^-^), 120 mM NMDG^+^/Glu^-^ (Glu^-^), 108 mM NMDG^+^+ 12 mM KVCH_3_SO_3_^-^ (CH_3_SO_3_^-^, K^+^) or 50% dilution of 120 mM NMDG7CH_3_SO_3_^-^ with isotonic sucrose (CH_3_SO_3_^-^, sucrose). The data were normalized with respect to the corresponding maximum value at the CH_3_SO_3_^-^ condition. Error bars are standard deviation: D129N, *n* = 4 for all experiments; D129L, *n* = 6 for all experiments; D129V, *n* = 18, 5, 5, 3 and 3 for CH_3_SO_3_^-^, Cl^-^, Glu^-^, CH_3_SO_3_^-^, K^+^ and CH_3_SO_3_^-^, sucrose, respectively.

### Neutralized D136 and D151 are impermeable to ions and protons

To evaluate other potential ion conducting phenotypes, two other countercharges, D136 and D151, located at the extracellular entrance of the gating pore water crevices (Fig. 1B), were mutated to alanine and asparagine. In contrast to D129 mutants, neutralization of D136 and D151 did not result in omega currents in NMDG^+^/CH_3_SO_3_^-^ under acidic, neutral or basic conditions (Fig. 4, A and B and fig. S4). However, their activation kinetics and voltage sensitivity were modulated by the external pH as with WT and the D129 mutants. Particularly, an apparent positive shift of the *Q-V* curve for the D136A, D136N and D151N mutants was observed at low pH. WE have shown previously that adding of 100 μM Cd^2+^ does not increase the net OFF charge *Q* in the V219C/D136A and V219C/D151A mutants of Ci-VSP (*48*), further confirms that neutralization of countercharge residues D136 and D151 cannot conduct omega currents.

**Fig. 4.**
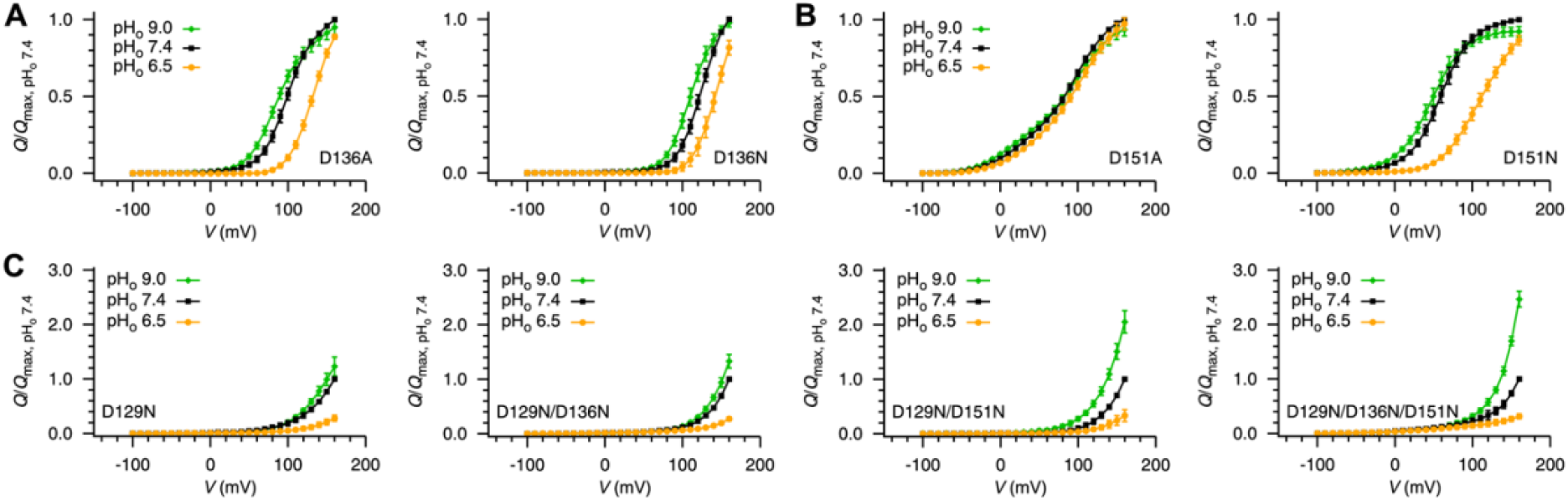
Single mutants of countercharge residues D136 and D151 are not ion permeable. (**A** to **C**) Normalized *Q*-*V* curves for the D136A and D136N (A), D151A and D151N (B), and D129N, D129N/D136N, D129N/D151N and D129N/D136N/D151N (C) mutants of Ci-VSP at different pHo with external recording buffer of 120 mM NMDG^+^/CH_3_SO_3_^-^ (CH_3_SO_3_^-^). The data were normalized with respect to the corresponding maximum value at pH_o_ 7.4. Error bars are standard deviation (D136A, *n* = 6, 8, 7 for pH_o_ 6.5, pH_o_ 7.4, pH_o_ 9.0, respectively; D136N, *n* = 7, 8, 7 for pH_o_ 6.5, pH_o_ 7.4, pH_o_ 9.0, respectively; D151A, *n* = 3, 8, 4 for pH_o_ 6.5, pH_o_ 7.4, pH_o_ 9.0, respectively; D151N, *n* = 6, 9, 5 for pH_o_ 6.5, pH_o_ 7.4, pH_o_ 9.0, respectively; D129N, *n* = 7, 10, 7 for pH_o_ 6.5, pH_o_ 7.4, pH_o_ 9.0, respectively; D129N/D136N, *n* = 8, 8, 8 for pH_o_ 6.5, pH_o_ 7.4, pH_o_ 9.0, respectively; D129N/D151N, *n* = 6, 8, 8 for pH_o_ 6.5, pH_o_ 7.4, pH_o_ 9.0, respectively; D129N/D136N/D151N, *n* = 6, 6, 6 for pH_o_ 6.5, pH_o_ 7.4, pH_o_ 9.0, respectively;).

Although countercharge residues D136 and D151 are not directly involved in the conduction of omega currents, they are still able to regulate the anionic omega current in the D129 mutants, likely by further changing the voltage sensitivity of gating (Fig. 4C). The asparagine substitution of D136 and/or D151 on the base of the D129N mutant amplifies the pH effect on the magnitude of the omega current. For example, the net charge *Q* of the triple mutant D129N/D136N/D151N increased about 2.5-fold at +160 mV when the pH changed from 7.4 to 9.0, while it is only about a 1.2-fold change for the single mutant D129N at the same condition.

### A molecular mechanism for CiVSD state dependency and ion selectivity

Formation of a continuous water pathway connecting the extracellular and intracellular solutions is a prerequisite for VSD ion conduction. This takes place only when the mutated gating charge residue(s) are close to the hydrophobic gasket region, leading to the state dependency of the omega currents (*32*, *34*–*36*, *43*, *48*). However, countercharges located in the S1-S3 segments of VSDs, have been associated to limited conformational rearrangement during voltage gating (*21*, *61*–*64*). To understand the molecular mechanism of the state dependent and selective conduction of omega currents, we performed MD simulations of the D129A and R226A mutants in Ci-VSD, as both mutants conduct transient proton currents during deactivation (*48*).

First, we examined gating pore water occupancy for the two mutants in the four functional states of Ci-VSD: “Down-minus”, “Down”, “Up” and “Up-plus”(*48*). We found evidence for a continuous water pathway through the entire gating pore of D129A and R226A, in both cases as they populate the “Down” state. However, the gating pore’s intra- and extracellular crevices appear disconnected at the hydrophobic gasket in the “down-minus” and “up-plus” states of D129A, and at the “down-minus”, “Up” and “up-plus” states of the R226A (fig. S5). In the “Up” state of the D129A mutant, water can hydrate the center hydrophobic gasket region but appears to be blocked at the site of D129A (fig. S5). We also quantitatively evaluated the propensity of the D129A gating pore to form a continuous water wire (Fig. 5A). Accordingly, R226A is able to form continuous water wires only in the “Down” state, while the D129A mutant can form continuous water wires in both its “Up” and “Down” states (but with a much reduced propensity).

**Fig. 5.**
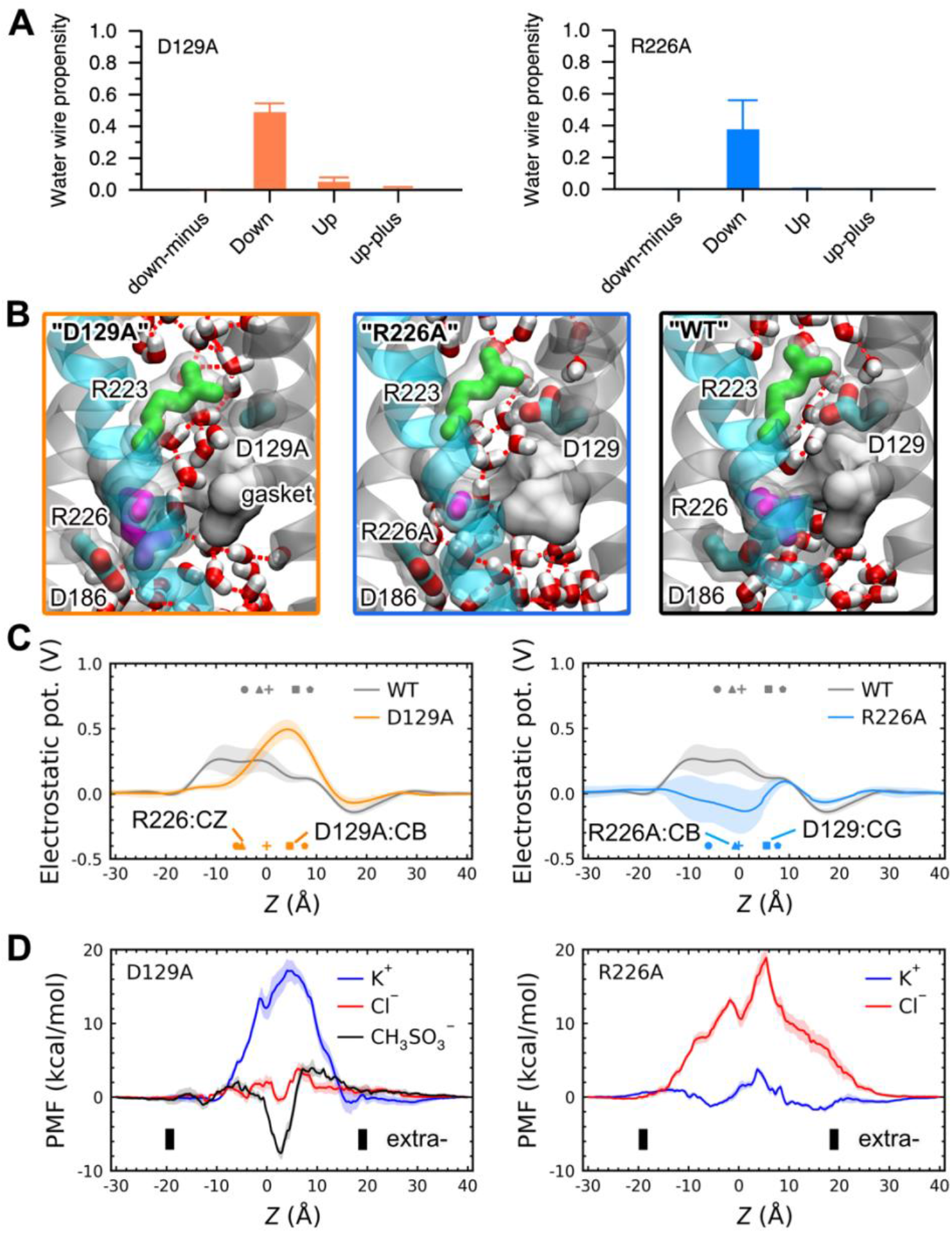
State dependent selective ion conduction through Ci-VSP mutants. (**A**) Propensity of forming a continues water wire connecting both sides of the D129A and R226A mutants at four major functional states. Error bars are standard deviation (*n* = 4). (**B**) Snapshots of the “Down” state D129A and R226A mutants and WT Ci-VSP from MD simulations, showing a continues hydrogen-bonded water wire connecting the two water crevices in the VSD of the two mutants. For clarity, hydrogen bonds between water molecules are highlighted in dashed lines, and the hydrophobic gasket is shown in surface representation. (**C**) Time-average electrostatic potential along a vertical line passing through the center of the WT and mutants of Ci-VSD. Transparent continuous errors are standard deviation. The average *z* coordinates of R223:CZ (pentagon), D129:CG/D129A:CB (square), center-of-mass of side-chain atoms of the hydrophobic gasket residues (plus), R226:CZ/R226A:CB (triangle) and D186:CG (circle) are shown in different markers. Error bars of the average *z* coordinates are too small to show. (**D**) One dimensional PMFs for ion conduction through the VSD of the D129A and R226A mutants Ci-VSP at the “Down” state. Transparent continuous errors are standard deviation. The black bars represent the approximate position of the membrane bilayer with the extracellular side (extra-) on the right.

Analysis of the MD trajectories showed that both, R226 side-chain occupancy in the hydrophobic gasket region and the rotameric flipping of R223 (in a salt-bridge with D129), prevent the formation of the continuous water wire through the “Down” state WT VSD (Fig. 5B). Neutralizing D129 and R226 enables the hydration of the entire gating pore by creating a leaking cavity at the hydrophobic gasket region, directly in R226A while indirectly in D129A, which allows R226 to reorient inwardly to interact with D186 (Fig. 5, B and C).

Electrostatic potential calculations show that neutralizing D129 makes the gating pore surface more positive than in WT, while neutralization of the gating charge R226 creates a negative electrostatic surface (Fig. 5C and fig. S6). We suggest that these electrostatic landscape changes help define the ion selectivity of the gating pore (*65*). One-dimensional potential of mean force (PMF) calculations were carried out to study the free energy for ion conduction through the D129A and R226A mutants. Consistent with our experimental observations, the energy barrier is about 4 kcal/mol for Cl^-^ and K^+^ ions to permeate in the D129A and R226A mutants, respectively.

But in the D129A mutant K^+^ encounters an energy barrier of over 16 kcal/mol to pass through the gating pore, while the cost of Cl^-^ permeation is about 20 kcal/mol in the R226A mutant (Fig. 5D). These results are abel to fully explain the different ion selectivities in the two mutants. The large anion CH_3_SO_3_^-^ experiences a deep energy basin of about −8 kcal/mol at the extracellular side of the hydrophobic gasket region, while the highest energy barrier is close to that for Cl^-^, which helps explain the lower permeability of CH_3_SO_3_^-^ relative to Cl^-^.

## Discussion

Voltage-sensing domains are conserved structural and functional modules, converting transmembrane potential changes into defined conformational rearrangements of its upstream or downstream effectors. A narrow hydrophobic “gasket”or charge transfer center at the core of most VSDs focuses the electric field into a narrow region and catalyzes the sequential and reversible translocation of positive gating charge residues across the electric field while preventing the permeation of physiological ions (*24*–*27*, *32*). However, under certain conditions gating pore “omega” currents have been observed in the VSDs of voltage-gated ion channels containing site-directed as well as naturally occurring mutations of S4 gating charge residues. These cation selective omega currents in sodium and calcium channels have been linked to certain channelopathies (*39*, *50*, *66*). In the present study, we report the discovery of a state dependent, anion selective omega current in Ci-VSP associated to single mutations in countercharge residues.

Just as with gating charge residue mutants, neutralization of the D129 countercharge in Ci-VSD can also create a continuous water permeation path (linked to its “Down” state) by changing the side-chain orientation of the gating charge residue occupying the hydrophobic gasket region. Gating charge residues in VSDs make extensive electrostatic interactions with countercharge residues, polar residues, as well as lipid headgroups during voltage gating (*19*, *53*, *67*–*69*). Neutralization of countercharge residues modifies this complicated network of interactions and flips the side-chain orientation of the nearby gating charge. This, in turn, opens a transient aqueous portal at the hydrophobic gasket region allowing ion conduction. In contrast, S4 translocation brings mutated gating charge residues across the electric field (i.e in histidine substitutions) or to the hydrophobic gasket region, making a continuous water pathway directly, and leading to proton and/or cation currents. Neutralization of gating charges and countercharges modifies the electrostatic landscape of the gating pore of VSDs, in a way that defines gating pore selectivity, cationic or anionic, respectively. In addition, the hourglass like morphology of the gating pore poses additional steric constrains that drive selectivity towards smaller ions, e.g., the small physiological ions K^+^, Na^+^, Cl^-^ have a higher permeability than the large organic ions guanidinium, NMDG^+^, CH_3_SO_3_^-^.

Neutralizing countercharge residues also changes the voltage sensitivity of gating of VSDs. The half-maximum activation potential of D129A shifts by over 100 mV in the positive direction compared with the WT. This increases the population of the anion conducting conformation (the “Down” state), upon activation, as much larger voltages are required to trigger the subsequent movement of the VSD from the “Down” state to more activated but less permeable conformations (e.g., the “UP” and “up-plus” states). Positive shift of the *Q*-*V* curve enhance the effects of pH on gating voltage sensitivity, by changing activation kinetics. High external pH accelerates the transition of the VSD from the resting “down-minus” state to the “Down” state, while increasing the”Down” state population at the same depolarization potentials. As expected, immobilization of S4 from deactivation by an engineered Cd^2+^ bridge leads to an increase in the dwell time of the “Down” state, which in turn increases the OFF currents (outward permeation of anions) during deactivation.

These findings define the state dependent anionic currents via VSDs countercharge mutants. While the underlying molecular mechanism here proposed extends our understanding of the functional, physiological, and pathological roles of countercharge residues in VSD containing membrane proteins. We expect that the present framework will help us uncover some of the unique mechanisms underlying channelopathies caused by countercharge mutations (*53*), develop new drugs targeting these diseases and design novel engineered proteins for scientific, industrial, and therapeutic applications.

## Materials and Methods

### Molecular biology and electrophysiological recordings

The cDNA of Ci-VSP with the catalytic center mutation (C363S) was subcloned into the pSP64T vector. This construct has no enzymatic activity but displays similar ON and OFF gating currents as the original one, and it has been used in the study of voltage sensitivity of Ci-VSP (*5*, *48*, *58*) and referred to as the wild type (WT) hereafter. The point mutations were generated using site-directed mutagenesis based on this construct. For multiple mutants containing a cysteine substitution, the background cysteine on the VSD (C159) was mutated to a serine to prevent the formation of an internal disulfide bond. Plasmids purified from miniprep were linearized and transcribed *in vitro* using the mMessage mMachine SP6 transcription kit (Ambion, Invitrogen). The mRNA was then diluted in RNase-free water to a final concentration of approximately 1 μg/μL. 50 nL of mRNA was injected into each freshly isolated *Xenopus laevis* oocyte, which was incubated in the standard oocyte saline (SOS) solution for 16-24 hours at 18 °C before recording.

Electrophysiological recordings were performed using a cut-open oocyte voltage-clamp setup (*56*, *57*). Currents were filtered at 10 kHZ using a low pass four-pole Bessel filter within the CA-1 amplifier (Dagan Corporation; Minneapolis, MN). A holding potential of −60 mV was applied, together with a 200 ms pre-pulse to −100 mV to maximally deactivate the WT and mutants of Ci-VSP. Test pulses from 160 mV to −100 mV in −10 mV decrements were used to evoke ON currents, which stepped back to −100 mV to generate OFF currents. In-house software GPatch and Analysis, kindly provided by Prof. Francisco Bezanilla, were used for data acquisition and analysis, respectively.

The basic external solution contained 120 mM N-methyl-D-glucamine (NMDG), 2 mM Ca(OH)_2_, 0.5 mM EDTA and 10 mM buffering agent: HEPES for pH7.4, MES for pH6.5 and CHES for pH9.0. Methanesulfonate acid (CH_3_SO_3_^-^) was used to adjust the pH to the corresponding value. The internal solution contained 120 mM NMDG, 2 mM EGTA, 10 mM HEPES, and the pH was adjusted to 7.4 using CH_3_SO_3_^-^. To change the external anion, HCl or glutamate was used for pH adjustment. To modify the external cation, a fraction of NMDG (12 mM) was replaced with KOH (12 mM), while keeping other components the same. Cd^2+^ was diluted into the external solution without the 0.5 mM EDTA at designed concentration from a 100 mM CdCl2 stock solution. Freshly prepared DTT solution of 1~2 mM was added to the external solution to chelate the Cd^2+^ ions.

The time constant of activation was calculated using the equation *⊺*_ON_=(*A*_1_*⊺*_1_+ *A*_2_*⊺*_2_)/(*A*_1_+*A*_2_) (*58*), where *A*_1_, *A*_2_ and *⊺*_1_, *⊺*_2_ are the amplitudes and time constants for the first and second exponentials, respectively, used to fit the decay phase of ON gating currents using the in-house software Analysis. Each individual *Q-V* curve was fitted and normalized using the non-linear least-squares minimization and curve-fitting (lmfit) package in Python (https://lmfit.github.io/), based on the Boltzmann distribution, *Q*(*V*)=1/{1+exp[*ze*_0_(*V*-*V*_1/2_)/*k*_B_*T*]}, in which *z* is the apparent gating charge, *V*_1/2_ is the half-maximum activation voltage, *e*_0_ is the elementary charge, *k*_B_ is the Boltzmann constant and *T* is the absolute temperature in Kelvin (*30*). The normalized *Q*-*V* curves were then averaged and fitted to get the best-fit values and standard deviations for the parameters *z* and *V*_1/2_ using the maximum likelihood by the Monte-Carlo Markov Chain method in the lmfit package (*30*).

### Molecular dynamics simulations

The D129A and R226A mutants of Ci-VSD at four different states were generated with the program VMD (*70*), using the corresponding centroid structures of the WT from the previous MD simulations (*48*). The ions in the bulk solution were readjusted to make the final systems electrically neutral. Following 5,000 steps energy minimization, each system was initially equilibrated for 50 ns. The coordinates of the last frame were then used to launch four independent 50 ns simulations with different initial set of velocities in the NPT ensemble to study the water occupancy and water wire propensity, and an additional 50 ns simulation in the NVT ensemble to study the electrostatic potential landscape. Positional restraints were applied to the alpha carbon atoms of residues 149 and 170 to prevent the drift of the protein. And “harmonicWalls” restraints of the collective variables interface (Colvars) module (*71*) were used to prevent the ions from entering the water crevices. The MD simulations were performed using the program NAMD (*72*). The CHARMM36 force field was used for the protein, lipids and ions (*73*, *74*), and the TIP3P model was used for water (*75*). The Langevin dynamics and the Nose-Hoover Langevin piston method were employed to keep the temperature and pressure at 300 K and 1 atm, respectively (*76*, *77*). The van der Waals interactions were smoothing switched between 10 - 12 Å. The long-range electrostatic interactions were calculated using the particle mesh Ewald (PME) method with a grid density of at least 1/Å^3^ (*78*). The timestep was 2 fs.

Water molecules near the principal axis of the VSD (−12 Å < x < 12 Å and −12 Å < y < 12 Å) were used to calculate the water occupancy with an interval of 2.0 Å in all the three dimensions. A total of 500 snapshots from one 50 ns simulation trajectory were used to calculate the average water occupancy. The continuous water wire propensity was calculated using the breadth-first algorithm (*48*, *79*). Results from all four trajectories of each state of each mutant were used to calculate the mean value and the standard deviation.

### Electrostatic potential calculations

Electrostatic potentials were calculated using the PMEPot plugin of VMD (*80*). The 800 snapshots from the last 40 ns trajectory were used to calculate the three-dimensional (3D) time-average electrostatic potentials. The 2D and 1D electrostatic maps were extracted from the 3D map along a plane or a vertical line passing through the center of the VSD (*81*). For the 1D map, the 800 snapshots were divided into four blocks and performed electrostatic potential calculation individually. The 1D results from the four blocks were then averaged to get the mean value and the standard deviation.

### Free energy calculations using umbrella sampling simulations

The umbrella sampling simulations were performed to calculate the 1D PMF for ion permeation through the D129A and R226A mutants of Ci-VSD at the “Down” state (*82*–*85*). The reaction coordinate was defined as the projection of the distance between the center-of-mass of the alpha carbon atoms of the hydrophobic gasket residues (I126, F161 and I190) and the permeating ion K^+^ or Cl^-^ or the sulfur atom of CH_3_SO_3_^-^ along the *z* axis using the Colvars module (*71*).

The last frame from the 50 ns equilibration simulation trajectory was used to generate starting configurations for the umbrella sampling simulations by moving the conducting ion to the target positions or sampling window centers using the Colvars module in each 1 ns equilibration simulation. Then, 10 ns production simulation was performed for each sampling window. A total of 141 windows (−30 Å < z < 40 Å, with an increment of 0.5 Å) were used for each PMF with a force constant of 10.0 kcal/mol/Å^2^ for the harmonic restraint. The sampling data were unbiased and combined using the weighted histogram analysis method (WHAM) to calculate the PMF (*86*). The last 8 ns sampling data were evenly divided into four blocks to get four 1D PMFs. The calculated PMFs were corrected first to remove the free energy offset between the two ends bulk water regions (*87*) and then averaged to get the mean and standard deviation values.

## Acknowledgments

We thank the Francisco Bezanilla laboratory at the University of Chicago for providing oocytes. We thank Drs. Tian Li, Michael David Clark and the members of the Perozo lab and the Roux lab for helpful advice and discussions.

## Author contributions

RS, BR, EP designed the whole study and analyzed the data. RS performed the electrophysiological experiments and MD simulations. RS wrote the initial manuscript. RS, BR, EP revised the manuscript.

## Data and materials availability

All data are available in the main text or the supplementary materials.

