## Supplemental Figures for "State Dependent Anionic Pore Currents Conducted by Single Countercharge Mutants in a Voltage-Sensing Phosphatase"

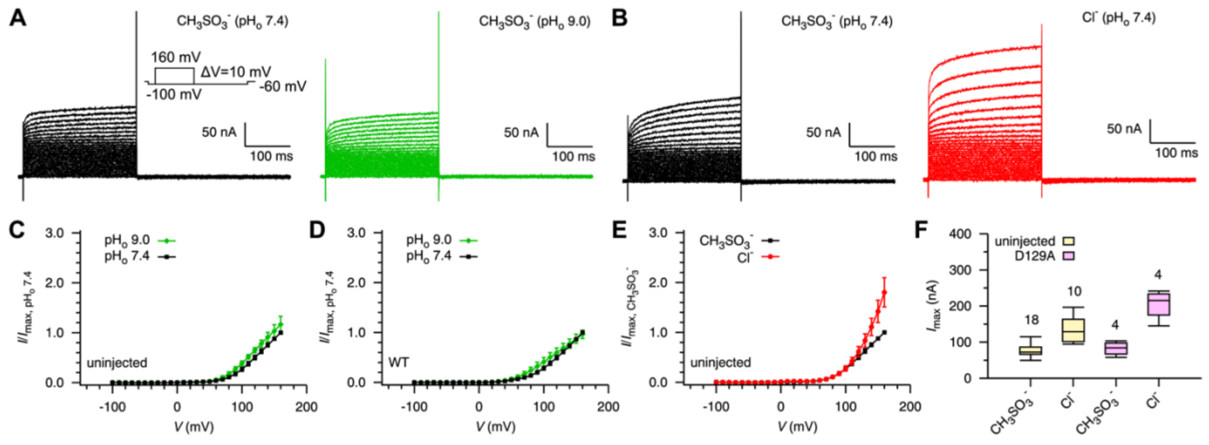


**Fig. S1. Electrophysiological characterization of un-injected Xenopus oocytes and oocytes injected with cRNA encoding the WT and D129A mutant of Ci-VSP.** (**A** and **B**) Representative currents of non-injected oocytes evoked at different pH values ($pH_o$) (A) or different components (B) of the external recording buffer. $CH_3SO_3^-$ and $Cl^-$ represent 120 mM $NMDG^+/CH_3SO_3^-$ and 120 mM $NMDG^+/Cl^-$ recording buffers, respectively. Inset is the voltage pulse protocol used to elicit the currents. (**C** and **D**) Normalized *I*-*V* curves for non-injected oocytes (C) and oocytes expressing WT Ci-VSP (D) with respect to the corresponding maximum value at $pH_o$ 7.4, where *I* is the magnitude of ON current at the end of each test pulse after linear leak subtraction. Error bars denote standard deviation (non-injected: $n$ = 7, 5 for $pH_o$ 7.4 and $pH_o$ 9.0, respectively; WT: $n$ = 7, 4 for $pH_o$ 7.4 and $pH_o$ 9.0, respectively). (**E**) Normalized *I*-*V* curves for non-injected oocytes with respect to the corresponding maximum value at the $CH_3SO_3^-$ buffer. Error bars denote standard deviation ($n$ = 18, 10 for $CH_3SO_3^-$ and $Cl^-$, respectively). (**F**) Comparison of the maximum *I* of non-injected oocytes and oocytes expressing D129A mutant of Ci-VSP. The data are displayed in box plots with the $n$ values being listed on the top of each box.

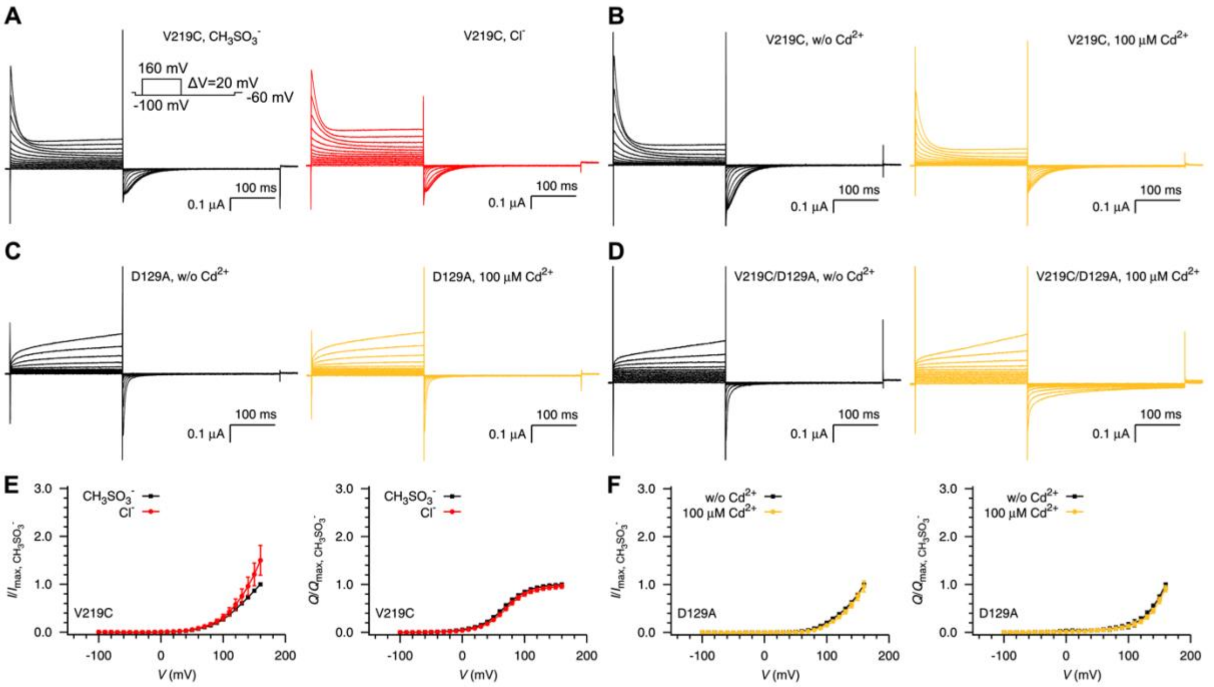


**Fig. S2. Effect of buffer components and $Cd^{2+}$ ions on the ON and OFF currents of Ci-VSP mutants.** (**A** to **D**) Representative currents of the V219C mutant with the external $CH_3SO_3^-$ or $Cl^-$ recording buffer (A); the V219C (B), D129A (C) and V219C/D129A (D) mutants in the absence of (w/o) or in the presence of 100 μM $Cd^{2+}$ in the external $CH_3SO_3^-$ buffer. Inset in (A) is the voltage pulse protocol used to elicit the currents. (**E**) Normalized *I*-*V* (left) and *Q*-*V* (right) curves for the V219C mutant. The data are normalized with respect to the corresponding maximum value at the $CH_3SO_3^-$ condition. Error bars are standard deviation ($n$ = 6). (**F**) Normalized *I*-*V* (left) and *Q*-*V* (right) curves for the D129A mutant. The data are normalized with respect to the corresponding maximum value at the w/o $Cd^{2+}$ condition. Error bars are standard deviation ($n$ = 4).

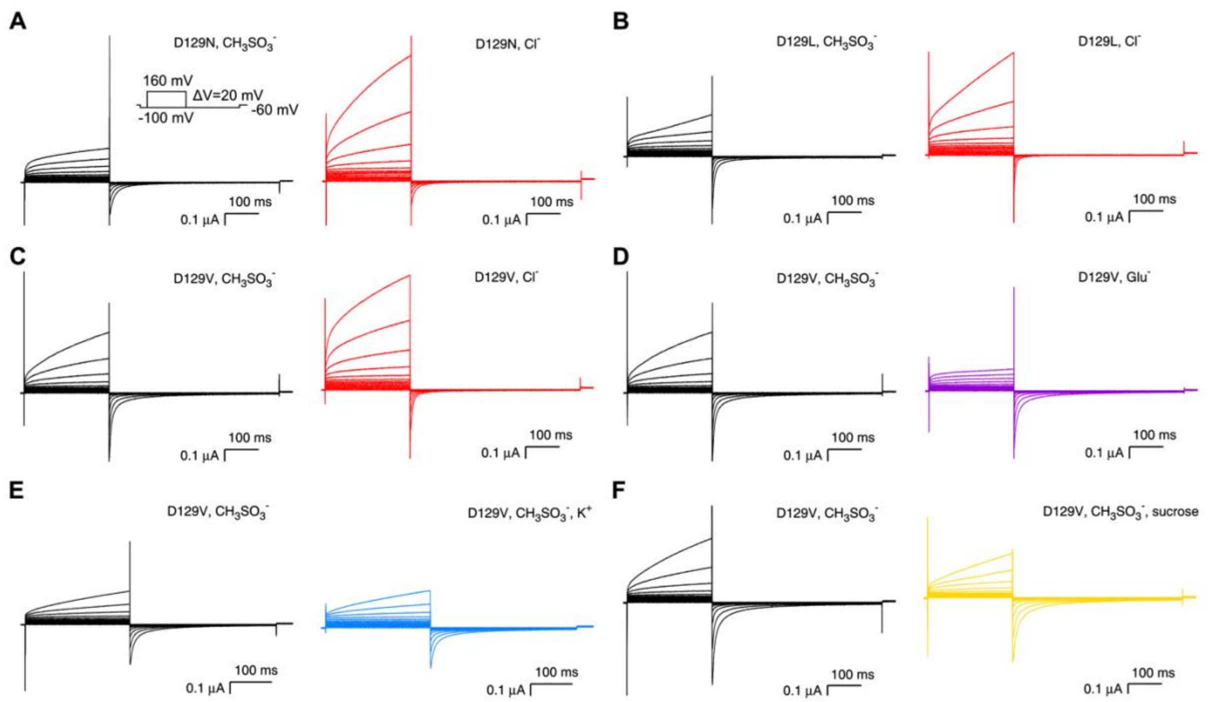


**Fig. S3. Anion conduction through single D129 mutants of Ci-VSP.** (**A**) Representative currents of D129N with external recording buffer of $CH_3SO_3^-$ (left) or $Cl^-$ (right). (**B**) Representative currents of D129L with external recording buffer of $CH_3SO_3^-$ (left) or $Cl^-$ (right). (**C** to **F**) Representative currents of D129V with external recording buffer of $CH_3SO_3^-$ (left) or other buffers of $Cl^-$ (right) (C), $Glu^-$ (right) (D), $CH_3SO_3^-$, $K^+$ (right) (E) and $CH_3SO_3^-$, sucrose (right) (F). Inset in (A) is the voltage pulse protocol used to elicit the currents.

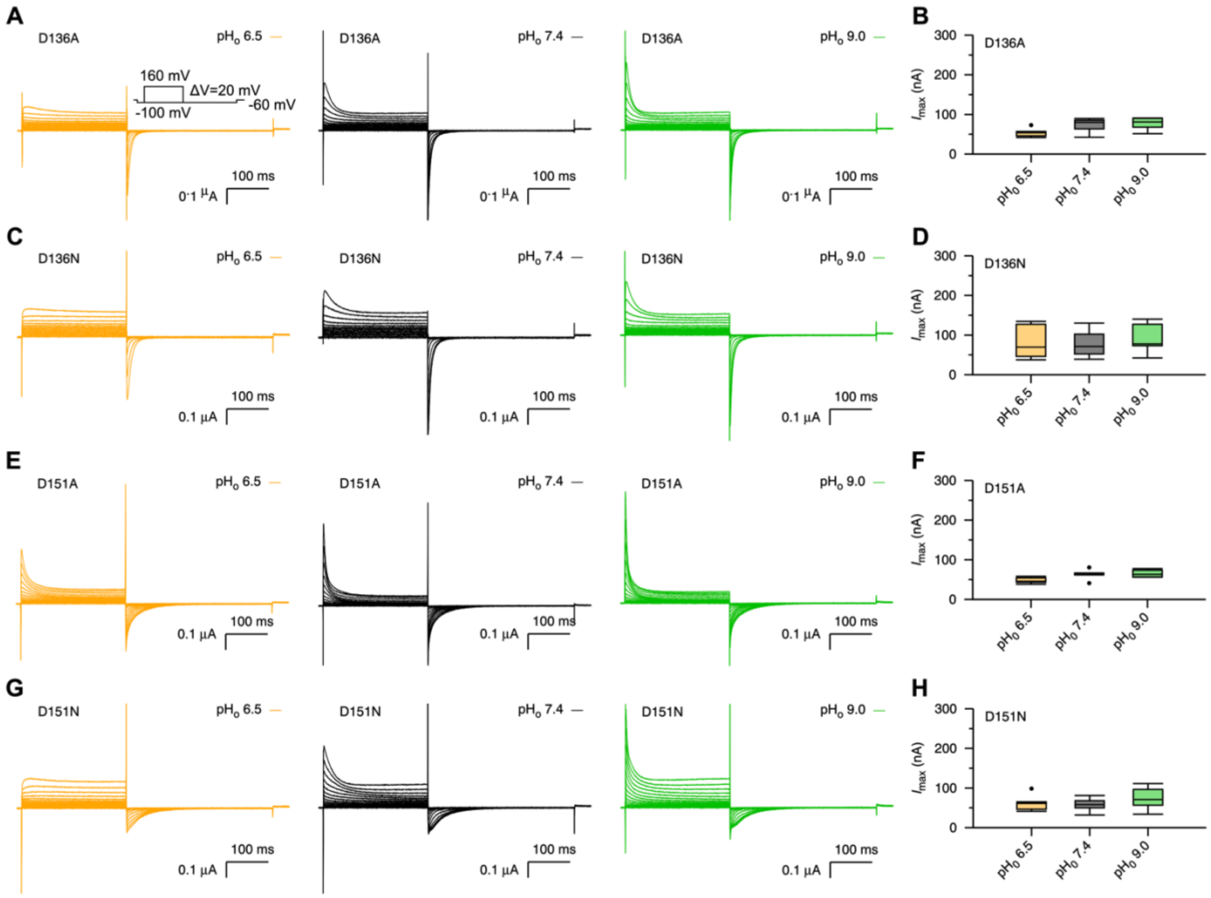


**Fig. S4. Effect of external pH on ON and OFF currents of single mutants of D136 and D151.** (**A, C, E** and **G**) Representative currents of D136A (A), D136N (C), D151A (E) and D151N (G) with external recording buffer of $CH_3SO_3^-$ at different external pH values. (**B**, **D**, **F** and **H**) Box plots of the maximum *I* of D136A (B), D136N (D), D151A (F) and D151N (H) with external recording buffer of $CH_3SO_3^-$ at different external pH values (D136A, *n* = 6, 8, 7 for pH$_o$ 6.5, pH$_o$ 7.4, pH$_o$ 9.0, respectively; D136N, *n* = 7, 8, 7 for pH$_o$ 6.5, pH$_o$ 7.4, pH$_o$ 9.0, respectively; D151A, *n* = 3, 8, 4 for pH$_o$ 6.5, pH$_o$ 7.4, pH$_o$ 9.0, respectively; D151N, *n* = 6, 9, 5 for pH$_o$ 6.5, pH$_o$ 7.4, pH$_o$ 9.0, respectively).

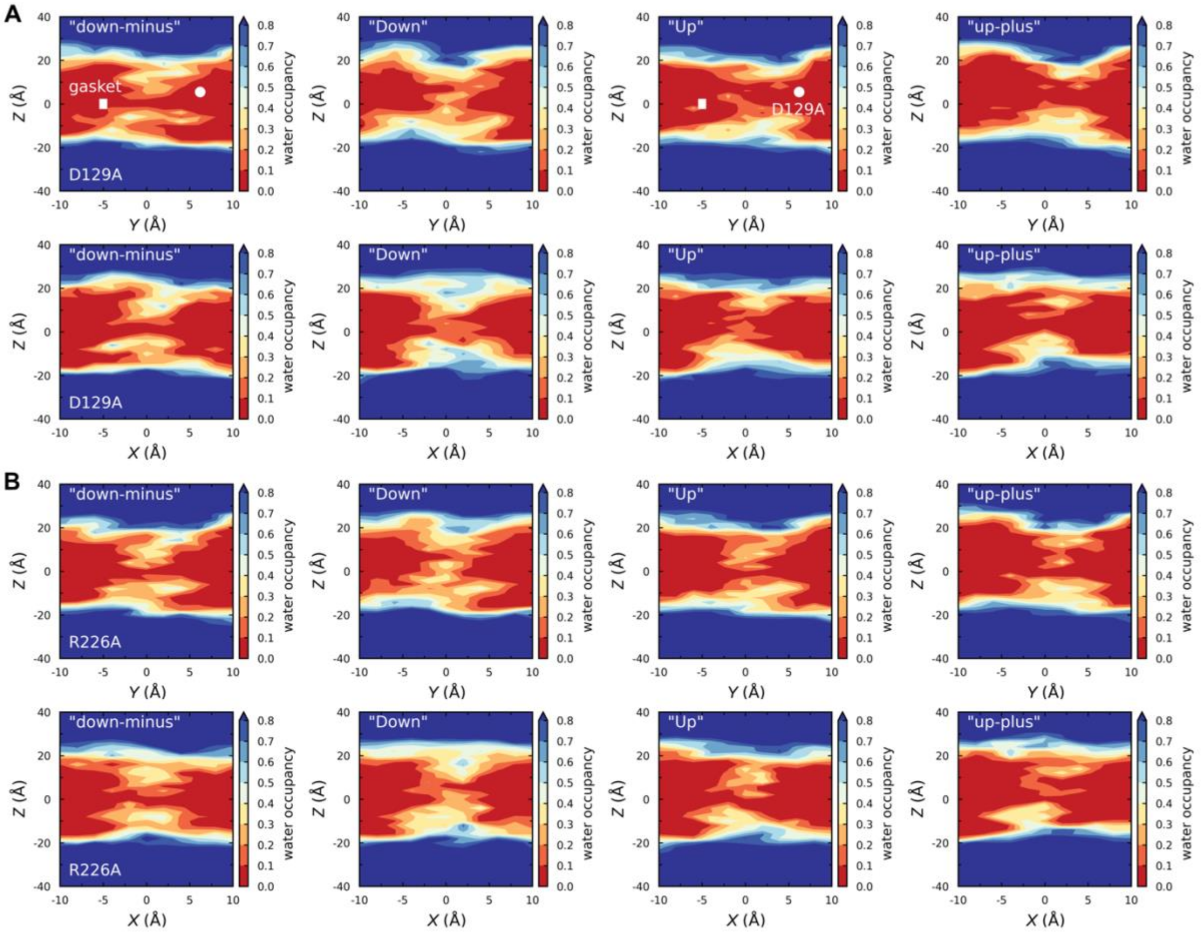


**Fig. S5. Continuous water pathway through the "Down" state VSD in D129A and R226A mutants of Ci-VSP.** (**A** and **B**) Projection of time- and space-average water occupancy in the *Y*-*Z* (top) and *X*-*Z* (bottom) planes thorough the extracellular and intracellular water crevices of VSD of the D129A (A) and R226A (B) mutants of Ci-VSP at four major functional states. The positions of the hydrophobic gasket and the countercharge residue D129 are shown with white bar and circle, respectively.

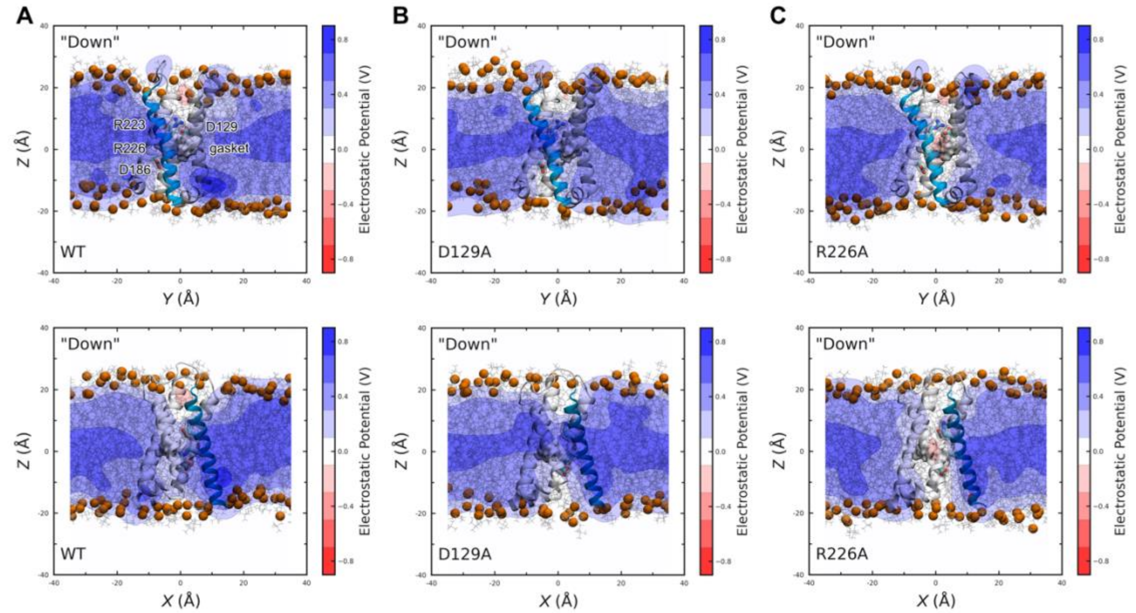


**Fig. S6. D129A and R226A mutations change the electrostatic potential landscape across the Ci-VSD.** (**A** to **C**) Two-dimensional time-average electrostatic potential maps passing through the center of the “Down” state VSD of WT (A), D129A (B) and R226A (C) mutants of Ci-VSP. The potentials are plotted in the *Y*-*Z* (top) and *X*-*Z* (bottom) planes. The S1-S4 transmembrane segments are shown in ribbon representation, with S4 being colored in cyan; the gasket residues in surface representation; and residues D129, R223, R226, D186 in stick and transparent surface representation. For clarity, only part of the lipids and their head group phosphate atoms are shown as grey lines and orange spheres, respectively.
